# Collider Scope: When selection bias can substantially influence observed associations

**DOI:** 10.1101/079707

**Authors:** Marcus R. Munafò, Kate Tilling, Amy E. Taylor, David M. Evans, George Davey Smith

**Author notes:** Corresponding author: Marcus R. Munafò, School of Experimental Psychology, University of Bristol, Bristol BS8 1TU, United Kingdom.; E.

## Abstract

Large-scale cross-sectional and cohort studies have transformed our understanding of the genetic and environmental determinants of health outcomes. However, the representativeness of these samples may be limited – either through selection into studies, or by attrition from studies over time. Here we explore the potential impact of this selection bias on results obtained from these studies, from the perspective that this amounts to conditioning on a collider (i.e., a form of collider bias). While it is acknowledged that selection bias will have a strong effect on representativeness and prevalence estimates, it is often assumed that it should not have a strong impact on estimates of associations. We argue that because selection can induce collider bias (which occurs when two variables independently influence a third variable, and that third variable is conditioned upon), selection can lead to substantially biased estimates of associations. In particular, selection related to phenotypes can bias associations with genetic variants associated with those phenotypes. In simulations, we show that even modest influences on selection into, or attrition from, a study can generate biased and potentially misleading estimates of both phenotypic and genotypic associations. Our results highlight the value of knowing which population your study sample is representative of. If the factors influencing selection and attrition are known, they can be adjusted for. For example, having DNA available on most participants in a birth cohort study offers the possibility of investigating the extent to which polygenic scores predict subsequent participation, which in turn would enable sensitivity analyses of the extent to which bias might distort estimates.

**Key Messages:** Selection bias (including selective attrition) may limit the representativeness of large-scale cross-sectional and cohort studies.

This selection bias may induce collider bias (which occurs when two variables independently influence a third variable, and that variable is conditioned upon).

This may lead to substantially biased estimates of associations, including of genetic associations, even when selection / attrition is relatively modest.

## Introduction

Understanding the impact of genetic and environmental factors on physical and mental health outcomes is critical if we are to develop effective preventive and treatment interventions. Large-scale cross-sectional and cohort studies provide an invaluable resource to support these efforts, in particular with respect to genetic influences, where the small effects associated with common genetic variants require very large samples to achieve adequate statistical power. A study can be used to draw conclusions about the population it represents, but generalizability to other populations depends upon us knowing exactly what the study population is. However, participants who volunteer to participate in studies may not be representative of the intended study population, in which case the actual study population is unknown (1).

Some studies may be relatively representative of the intended study population at inception through rigorous efforts to ensure representative recruitment (e.g., birth cohort studies). However, as they mature the likelihood is that attrition from the study will be non-random, so that the cohort becomes less representative of the intended population as time goes on. Alternatively, the reverse may be true –the study may be unrepresentative at inception, but with low attrition. Selection bias can also occur if a sub-set of participants within a study is selected for more detailed investigation (e.g., genotyping) on the basis of having most data available, or volunteering for further follow-up (2). There is already clear evidence from existing large-scale population studies that they are subject to a degree of selection bias. For example, higher genetic risk scores for schizophrenia are consistently associated with non-completion of questionnaires by study mothers and children, as well as non-attendance at data collection clinics, in the Avon Longitudinal Study of Parents and Children (ALSPAC) (3) (see Box 1).

#### Box 1. The Avon Longitudinal Study of Parents and Children.

Birth cohort studies are also not immune to problems of selection bias, where retention in the study may be related to a variety of participant characteristics. The Avon Longitudinal Study of Parents and Children (ALSPAC) recruited pregnant women living in the administrative county of Avon with expected delivery dates between 1st April 1991 and 31st December 1992. These women, their partners and their offspring have been followed up ever since via questionnaires and clinics. ALSPAC originally captured data on 14,541 pregnancies (75% of eligible women) (19, 36), but inevitably retention in subsequent data collection sweeps (postal questionnaires and clinic assessments) was less than 100%. We see that higher body mass index (BMI) is associated with lower odds of subsequent retention in both mothers (N = 11,319, OR per SD increase in BMI 0.85, 95% CI 0.81 to 0.88), for retention between 2008 and 2011 using pre-pregnancy BMI as a predictor, and offspring (N = 7,954, OR 0.91, 95% CI 0.87 to 0.96), for retention at age 18 using BMI at age 7 as a predictor. Similarly, among smoking mothers in ALSPAC, heaviness of smoking is associated with lower odds of retention (N = 3,534, OR per additional cigarette smoked per day just prior to pregnancy 0.97, 95% CI 0.96 to 0.98). If low BMI and maternal non-smoking are both related to continuing participation in ALSPAC, this would tend to lead to the association between BMI and maternal smoking being negatively biased (i.e., we would expect to see a more negative association between smoking and BMI in ALSPAC than in the true underlying population).

Attrition from cohort studies may result in biased estimates of socioeconomic inequalities, and the degree of bias may worsen as participation rates decrease (4). However, it is often argued that representativeness is not necessary in studies of this kind (5-9), although this is not universally accepted (10). In particular, for genetic variants, where conventional confounding is low (11), it has been argued, even by those concerned about selection bias, that any problems associated with a lack of representativeness may be modest (10, 12). Here we ask: What is the impact of selection bias on the results obtained from these studies? We take the perspective that selection bias can amount to conditioning on a collider (i.e., conditioning on a variable that is independently influenced by two other variables).

## Collider Bias

It is widely acknowledged that selection bias will distort prevalence estimates. This can be clearly seen in differences between participants at baseline and at subsequent assessments in cohort studies, such as when we compare the original ALSPAC sample with those who attended later clinics (see Box 1). It can also be seen in differences between a study sample and the source population from which it is drawn; for example, the UK Biobank study differs relative to the general population in the UK (see Box 2). However, it is often assumed that whilst selection bias will have a strong effect on prevalence estimates, it should not have a strong impact on observed associations between variables (8). This overlooks the fact that selection can induce collider bias (see Figure 1), which can lead to biased observational and genetic associations. This bias can be towards or away from any true association, and can distort a true association or a true lack of association.

#### Box 2. UK Biobank.

The UK Biobank is a cross sectional study, which recruited over 500,000 individuals aged between 40 and 69 years between 2006 and 2010 (see http://www.ukbiobank.ac.uk/). Individuals in this age group living within a 25 mile radius of any of the 22 assessment centres across the UK were identified from NHS patient registers (37). In total, around 9 million individuals were invited to participate. However, UK Biobank was only able to achieve a 5% response rate (~500,000 participants recruited from ~9,000,000 invited, personal communication, UK Biobank, 8th July 2016), and the resulting sample is not representative of the UK population as a whole. For example, the proportion of current smokers is relatively low in UK Biobank (19% in the general population vs 11% in UK Biobank, equivalent to an OR of 1.90) (38), as is the proportion with no qualifications (25% vs 17%, equivalent to an OR of 1.63) (39). Unsurprisingly, therefore, participants in UK Biobank have far lower rates of 5-year mortality than the UK population as a whole (40). Clearly, agreeing to take part in the UK Biobank study is associated with a number of characteristics that will reflect, for example, health status and social position. If nonsmoking and having qualifications are both causally related to participation in UK Biobank, we would expect the association between smoking and having qualifications to be positively biased (i.e., we would expect to see a more positive association between genetic variants positively associated with smoking and whether participants had educational qualifications in UK Biobank than in the true population). The problem is possibly compounded in genetic studies using the first release of genomewide association data in UK Biobank, which used two genotyping arrays, one of which was applied to a nested case-control study of smoking and lung function (UK BiLEVE) (41). The first release genetic data are therefore further subject to selection bias relative to UK Biobank as a whole (although this will no longer be the case when the full release of genomewide association data becomes available).

**Figure 1.**
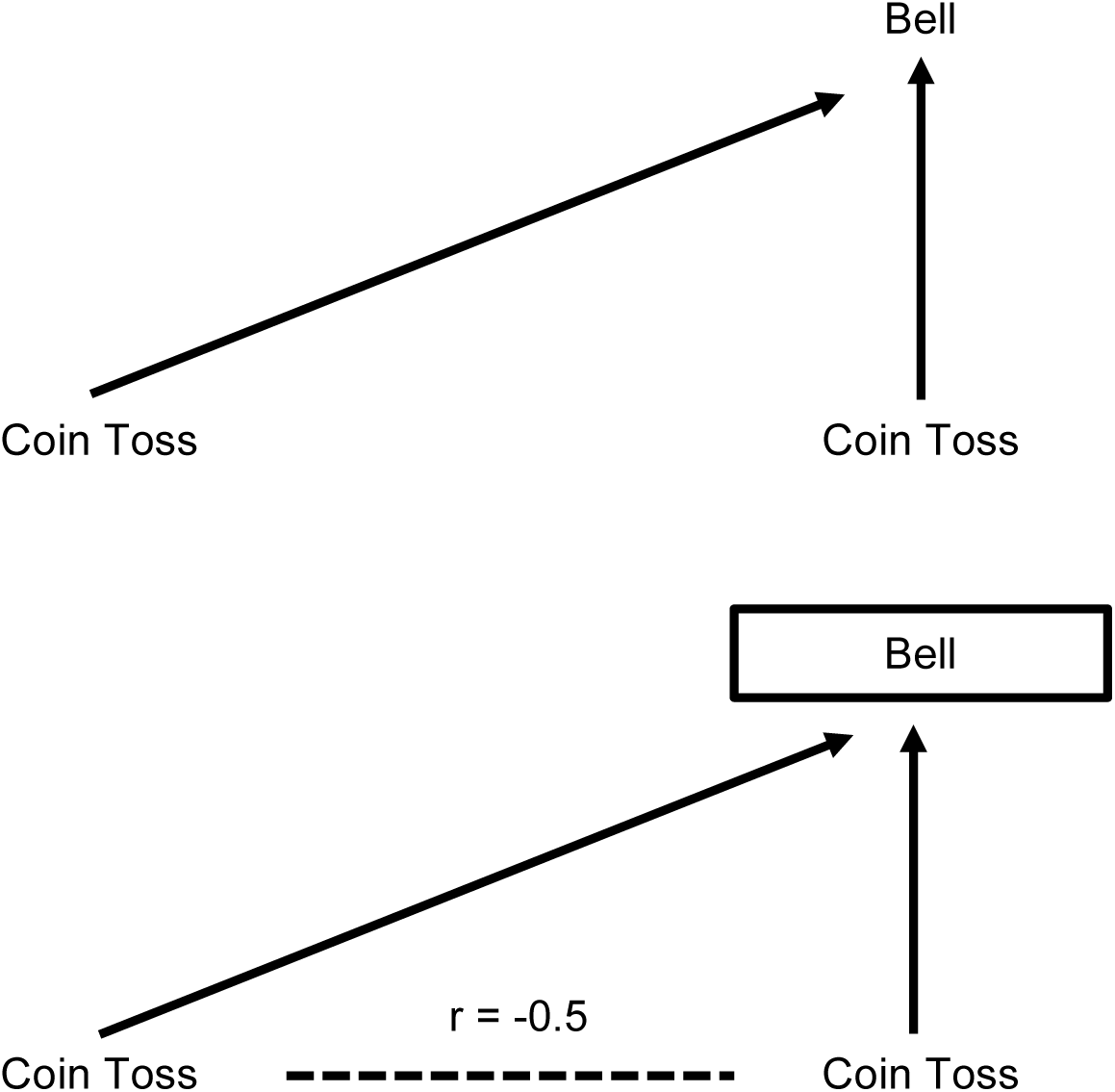
Illustration of collider bias. The basic premise of collider bias is shown. In this example, a bell is sounded whenever either coin come up ‘heads’. The result of one coin toss is independent of the other. However, if we hear the bell ring (i.e., we condition on the bell ringing), then if you see a tail on one coin you know there must be a head on the other – the two coin results are no longer independent and a spurious inverse correlation has been induced. Reproduced from Gage SH, Davey Smith G, Ware JJ, Flint J, Munafò MR (2016) G = E: What GWAS Can Tell Us about the Environment. PLoS Genet 12(2): e1005765. doi:10.1371/journal.pgen.1005765

Collider bias occurs when two variables (*X* and *Y*) independently cause a third variable (*Z*). In this situation, *Z* is a collider, and statistical adjustment for *Z* will bias the estimated causal association of *X* (exposure) on *Y* (outcome) (see Figure 2). Statistical adjustment of the *XY* association for a variable *Z* is equivalent to observing this association in a sub-population where all individuals share the same value of *Z* (1, 13). Hence if both *X* and *Y* cause participation in a study (*Z*), then investigating associations in the selected sample (i.e., with *Z* = 1, indicating participation in the study) is equivalent to conditioning on *Z*, which in turn may induce collider bias.

**Figure 2.**
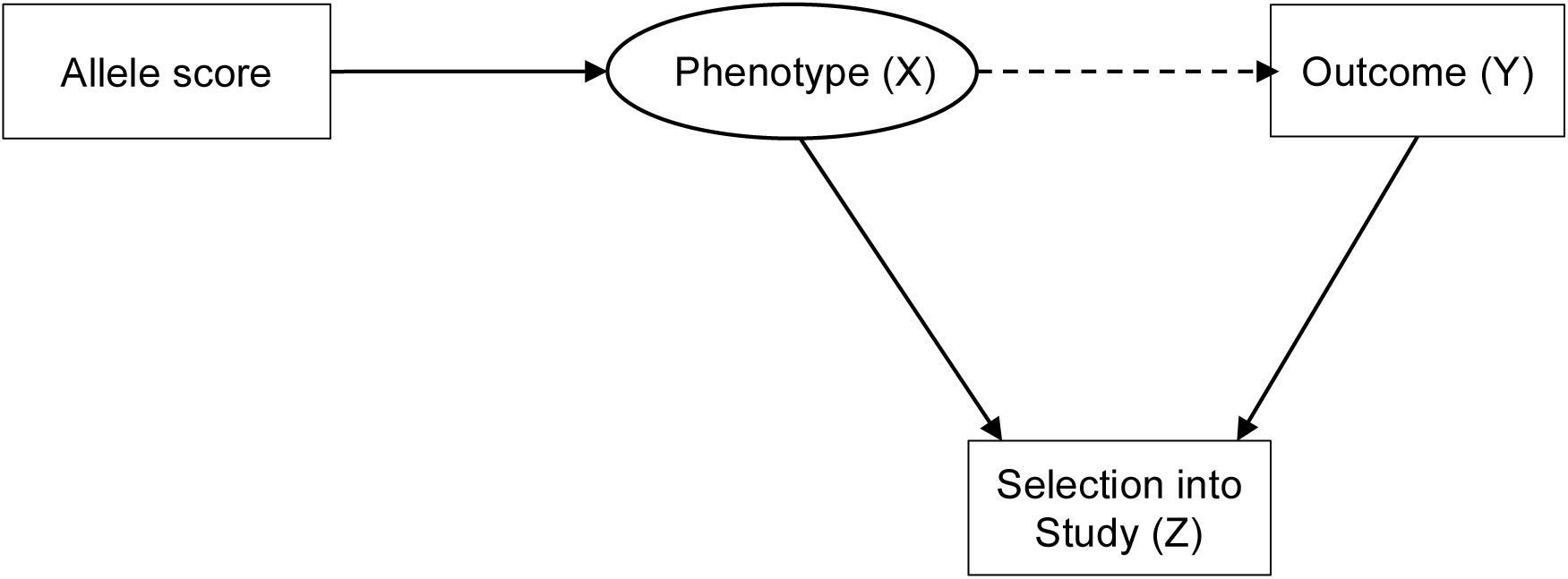
Illustration of selection bias simulation. In the entire population there is no association between allele score and outcome. Selection into the study (either through voluntary participation at baseline, or attrition over time) induces an association between allele score and outcome (collider bias).

In other words, sample selection can bias associations between variables that influence participation or retention in a study. This can include inducing spurious associations when no such association exists in the population from which the sample is drawn or, if two variables are correlated in the intended study population and both cause selection, biasing the estimated correlation in the selected sample. Moreover, this selection bias will apply to the genetic correlates (or other ancestors) of these variables, unless the phenotypes are also controlled for. Therefore if genes Gx and Gy cause *X* (exposure) and *Y* (outcome) respectively, and both *X* and *Y* influence participation, then in the selected sample Gx will appear to be associated with *Y* (unless *X* is also controlled for). More complex situations can also give rise to collider bias, such as when the outcome (*Y*) doesn’t directly *cause* selection into the study (i.e., it is a downstream consequence of something else that *is* causing selection into the study).

If two traits influence participation (and therefore contribute to selection), selection bias amounts to implicitly conditioning on their common effect (i.e., participation) (1, 14). This can in principle lead to biased associations between these two traits. There are exceptions to this depending on the distribution of the outcome and the parametric analysis model used. For example, if the outcome (*Y*) is a binary phenotype, and logistic regression is used, then the odds ratio for the association between the SNP and outcome may be unbiased even when the outcome causes selection (as is true of case-control studies) (15). We have previously argued that these effects may be greater in case-control studies than prospective studies, and that since genetic associations have been similar across study designs, the impact of selection bias may in fact be modest (12). We have also previously argued that because conventional confounding is typically low for single genetic variants, problems of selection bias will be less in this context (10). However, given the rapid growth in studies using data from highly selected samples such as UK Biobank, and the use of genetic risk scores rather than single genetic variants, we revisited this question, and used simulation to explore the potential impact of even relatively weak effects on participation. Given empirical evidence of selection in cross-sectional and cohort studies, what is the potential impact of this on observed phenotypic and genotypic associations?

## Simulations

We simulated data on an allele score, a phenotype and an outcome, where both the phenotype and outcome influence selection into the study, but there was no association between the allele score and the outcome in the underlying population (see Figure 2). The simulation scenario was based loosely on the UK Biobank, and we simulated selection into the study, so all the data on non-selected individuals are missing and therefore imputation is not a potential solution (see below), because this requires some data on which to base the imputation (16). All variables were Normally distributed, with standard deviation of 1, and the sample size of the underlying complete population was 9,000,000. We assumed that phenotype and outcome had independent effects (i.e., no interaction on the additive scale) on the odds of selection into the sample, and for convenience we set these effects to be equal, and examined a weak association (OR of 1.2 for missingness for a 1 SD increase in phenotype/outcome) and two stronger associations (ORs of 1.5 and 1.8). These odds ratios are similar to estimates of the likelihood of participation in UK Biobank for individuals with any educational or vocational qualifications and for non–smokers, respectively (see Box 2), and indicate a difference in mean phenotype/outcome of 0.2 SD, 0.4 SD and 0.6 SD between those participating and those not participating. We varied the correlation between the allele score and the phenotype (between r = 0.05 and r = 0.30) to simulate genetic instruments explaining between 0.25% and 9% of the variance in phenotypes. These values are in the typical range for the association between common genetic variants, or polygenic risk scores comprising multiple common variants, and complex phenotypes. For example, the rs16969968 variant accounts for approximately 1% of the phenotypic variance in cigarette consumption (17), while the polygenic risk score for height captures approximately 9% of phenotypic variance (18). We controlled the baseline risk of selection into the sample, resulting in a selected sample of approximately 500,000 people. The analysis was an unadjusted regression of outcome on allele score (i.e. not adjusting for the phenotype). We simulated a true null association (i.e. in the whole population, the regression coefficient for outcome on allele score is zero). We simulated each scenario 100 times. We then repeated the simulations with the addition of a causal effect of the phenotype on the outcome, with a regression coefficient of 0.1.

The results of this simulation study are shown in Table 1 (no causal effect of P on O) and Table 2 (causal effect of P on O). Where there is no causal effect of P on O, the effects of selection bias are strongest for stronger independent selection effects, and also where the allele score is more strongly associated with the phenotype (Table 1). However, even for moderate associations between missingness and both phenotype and outcome (OR = 1.5 for both phenotype and outcome) and between allele score and phenotype (r = 0.1, 1% variance explained by allele score) the confidence intervals contains zero only 89% of the time, and this continues to decrease with both greater strength of association between phenotype, outcome and missingness, and stronger association between allele score and phenotype.

**Table 1.**
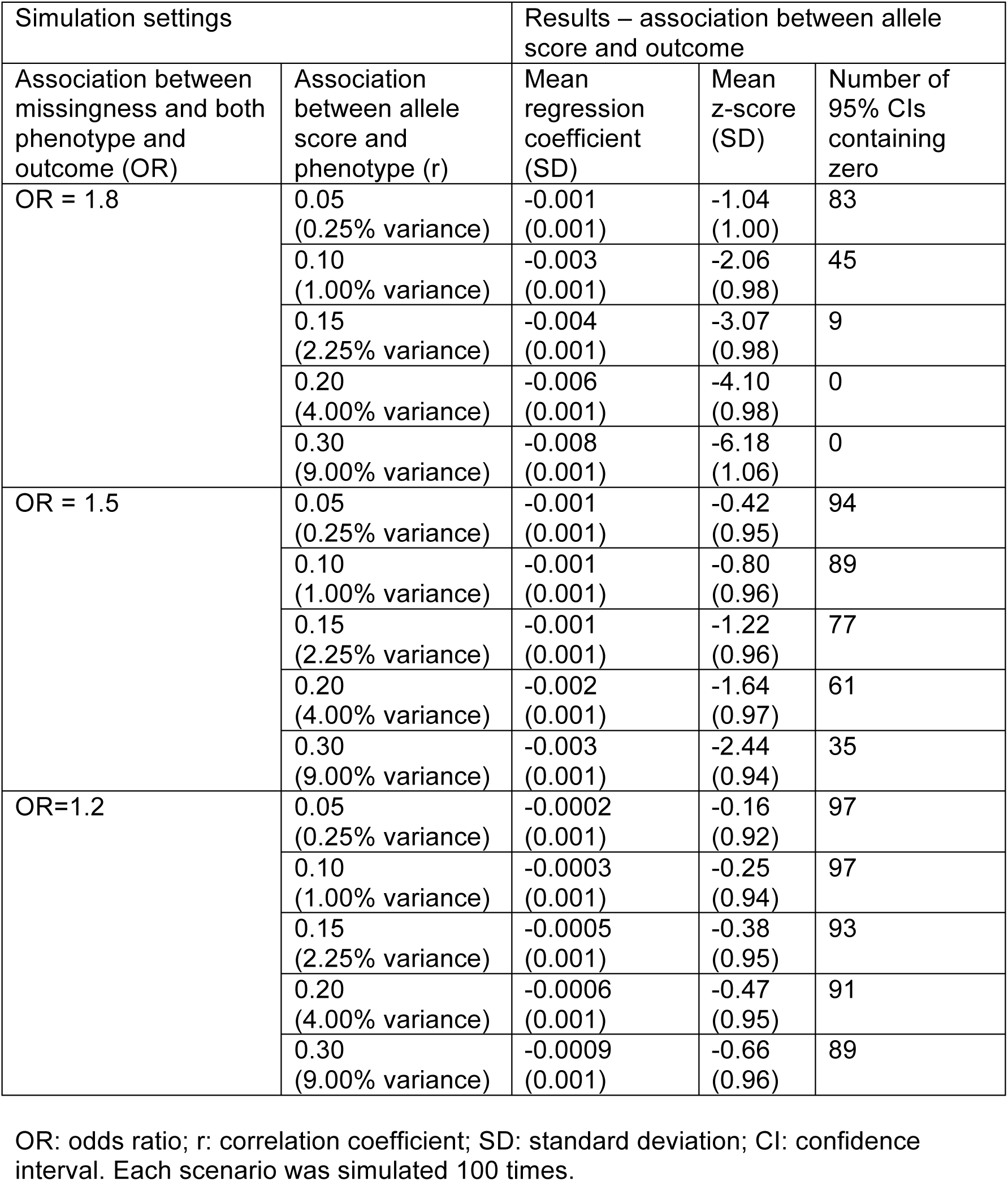
Results of simulation study showing the selection bias in estimating an association that is null in the underlying population.

| Simulation settings |  | Results – association between allele score and outcome |  |  |
| --- | --- | --- | --- | --- |
| Association between missingness and both phenotype and outcome (OR) | Association between allele score and phenotype (r) | Mean regression coefficient (SD) | Mean z-score (SD) | Number of 95% CIs containing zero |
| OR = 1.8 | 0.05<br>(0.25% variance) | -0.001<br>(0.001) | -1.04<br>(1.00) | 83 |
|  | 0.10<br>(1.00% variance) | -0.003<br>(0.001) | -2.06<br>(0.98) | 45 |
|  | 0.15<br>(2.25% variance) | -0.004<br>(0.001) | -3.07<br>(0.98) | 9 |
|  | 0.20<br>(4.00% variance) | -0.006<br>(0.001) | -4.10<br>(0.98) | 0 |
|  | 0.30<br>(9.00% variance) | -0.008<br>(0.001) | -6.18<br>(1.06) | 0 |
| OR = 1.5 | 0.05<br>(0.25% variance) | -0.001<br>(0.001) | -0.42<br>(0.95) | 94 |
|  | 0.10<br>(1.00% variance) | -0.001<br>(0.001) | -0.80<br>(0.96) | 89 |
|  | 0.15<br>(2.25% variance) | -0.001<br>(0.001) | -1.22<br>(0.96) | 77 |
|  | 0.20<br>(4.00% variance) | -0.002<br>(0.001) | -1.64<br>(0.97) | 61 |
|  | 0.30<br>(9.00% variance) | -0.003<br>(0.001) | -2.44<br>(0.94) | 35 |
| OR=1.2 | 0.05<br>(0.25% variance) | -0.0002<br>(0.001) | -0.16<br>(0.92) | 97 |
|  | 0.10<br>(1.00% variance) | -0.0003<br>(0.001) | -0.25<br>(0.94) | 97 |
|  | 0.15<br>(2.25% variance) | -0.0005<br>(0.001) | -0.38<br>(0.95) | 93 |
|  | 0.20<br>(4.00% variance) | -0.0006<br>(0.001) | -0.47<br>(0.95) | 91 |
|  | 0.30<br>(9.00% variance) | -0.0009<br>(0.001) | -0.66<br>(0.96) | 89 |

OR: odds ratio; r: correlation coefficient; SD: standard deviation; CI: confidence interval. Each scenario was simulated 100 times.

**Table 2.**
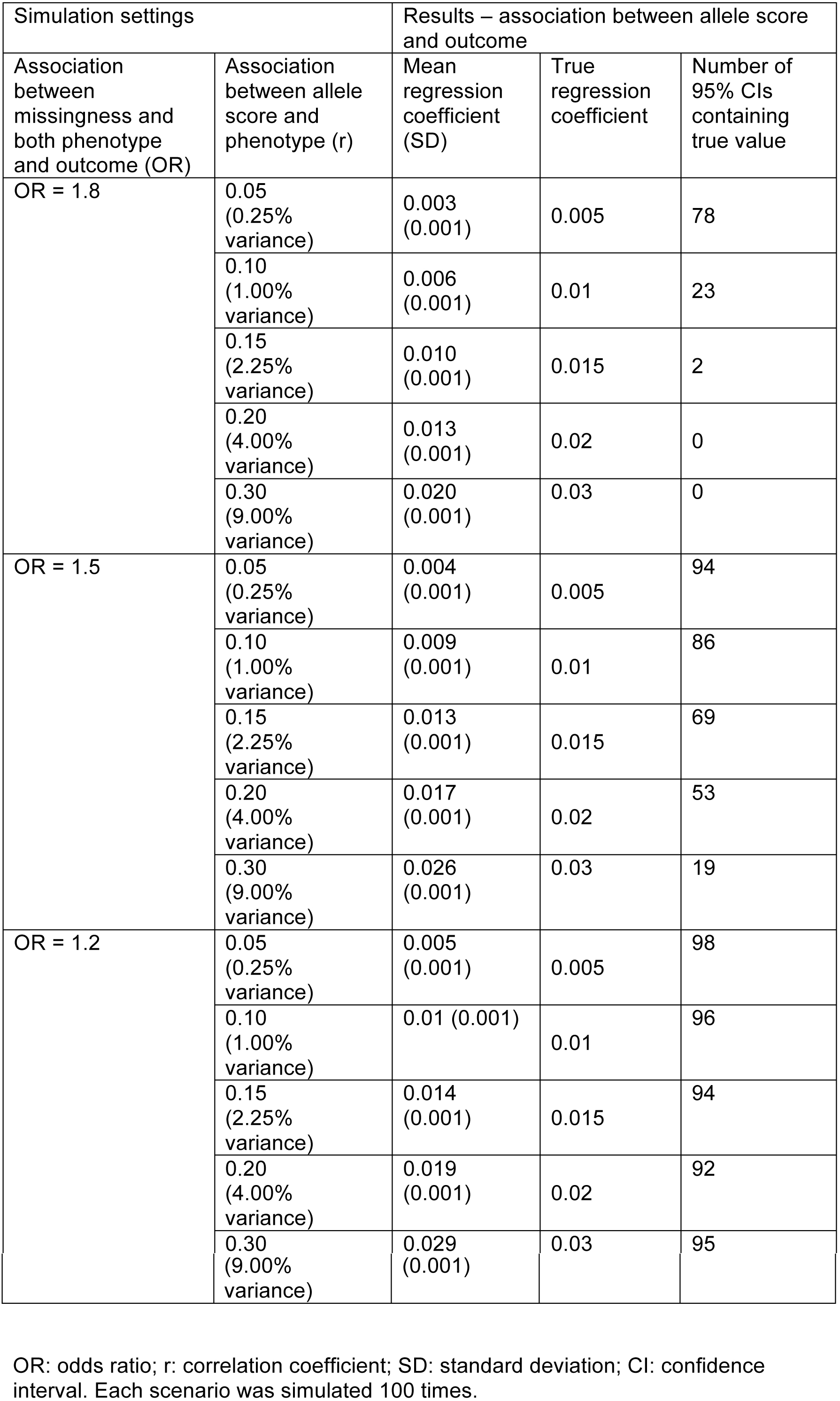
Results of simulation study showing the selection bias in estimating an association that is not null in the underlying population (regression coefficient for outcome on phenotype is 0.1)

Where there is a causal effect of P on O, the results are broadly similar, except that on the whole the confidence intervals had lower coverage than for the equivalent situation with no causal association.

We also explored associations between known risk factors and outcomes in a representative birth cohort and a selected sub-study. We used ALSPAC as the birth cohort. Initially 14,541 pregnant women who were expected to give birth between 1 April 1991 and 31 December 1992 were recruited into the study in the South West region of England (19). The study website contains details of all data available through a fully searchable data dictionary (http://www.bris.ac.uk/alspac/researchers/data-access/data-dictionary/). Ethics approval for the study was obtained from the ALSPAC Ethics and Law Committee and the Local Research Ethics Committees. We also used the Accessible Resource for Integrated Epigenomics Studies (ARIES), a sub-study of ALSPAC where a subset of 1,018 mother-offspring pairs were selected based on availability of DNA samples at two time points for the mother (at an antenatal clinic and at a follow-up clinic when their offspring were mean age 15.5 years) and three time points for the offspring (at birth, childhood, and adolescence (2). We investigated the association between a genetic risk score for smoking (ever vs never) and maternal education in ALSPAC, and in the ARIES sub-sample. The results are shown in Table 3, and indicated that the genetic risk score for smoking and maternal education are associated in ARIES, but not in the full sample.

**Table 3.** Associations between a genetic risk score for smoking and maternal education, in ALSPAC and ARIES.

| **Association between genetic risk score and ever smoking in ALSPAC** |  |  |  |
| --- | --- | --- | --- |
|  | N | OR (95% CI) | P |
| Smoking genetic risk score <sup>1</sup> | 7,291 | 1.07 (1.02 to 1.12) | 0.003 |
| **Association with being in the ARIES sub-study** |  |  |  |
|  | N | OR (95% CI) | P |
| Smoking (ever vs never) | 13,249 | 0.59 (0.52 to 0.68) | <0.001 |
| Smoking genetic risk score <sup>1</sup> | 7,837 | 1.00 (0.93 to 1.07) | 0.92 |
| Maternal education <sup>2</sup> | 12,493 | 1.86 (1.58 to 2.19) | <0.001 |
| **Association between smoking/smoking genetic risk score and maternal education in ALSPAC and ARIES** |  |  |  |
| **ALSPAC** |  |  |  |
|  | N | OR (95% CI) | P |
| Smoking (ever vs never) | 12,118 | 0.45 (0.40 to 0.50) | <0.001 |
| Smoking genetic risk score <sup>1</sup> | 7,046 | 1.01 (0.95 to 1.08) | 0.74 |
| **ARIES** |  |  |  |
|  | N | OR (95% CI) | P |
| Smoking (ever vs never) | 986 | 0.61 (0.44 to 0.84) | 0.003 |
| Smoking genetic risk score <sup>1</sup> | 791 | 1.20 (1.02 to 1.41) | 0.03 |

1. Genetic risk score including variants reaching $P < 0.05$ for association with ever vs never smoking in the Tobacco and Genetics Consortium GWAS (see Supplementary Material). Associations are per SD increase in genetic risk score. 2. Degree vs no degree.

## Conclusions

Our results indicate the potential for selection/attrition to generate biased and potentially misleading estimates of both phenotypic and genotypic associations. In particular, when polygenic scores (associated with a phenotype) that combine many genetic variants are used, association between the phenotype and participation will cause the score to be more strongly related to participation than each individual variant is. This, in turn, can potentially lead to serious bias. For this reason, studies using polygenic scores, genome-wide allelic scores (20), and/or whole-genome genetic correlations (21, 22) are most at risk of producing biased and potentially misleading results where there is reason to believe the study sample is not representative of the study population but the mechanism of selection is unknown.

The magnitude of effects we observed in our simulations, based on credible estimates of associations between both a phenotype or outcome and missingness, and between a polygenic score and a phenotype, are comparable with many reported associations derived from large but selected samples, such as between personality and cognitive function, and a range of physical and mental health outcomes (23, 24), and between chronotype (i.e., “morningness”) and years of education (25). Such associations could therefore plausibly be generated by selection bias. An appreciation of the potential impact of selection bias may also resolve inconsistencies in the literature, and help to explain apparently paradoxical findings. For example, genetic correlations between cognitive ability and a range of psychiatric disorders have been reported to differ in childhood and older age (26). One possible interpretation is that this is due to age-dependent pleiotropy, but another is that this is an artefact of different selection bias pressures at different ages. An example serves to illustrate this. Polygenic risk scores that maximally capture schizophrenia liability are associated with increased psychotic experiences in ALSPAC participants, but scores that use more stringent thresholds for including genetic variants are associated with *reduced* psychotic experiences (27). Since missing data are likely to be greater for participants who report psychotic experiences, as well as for those at higher genetic risk of a psychotic disorder, psychotic experiences may be relatively under-represented in participants with higher genetic risk, compared to those with lower genetic risk (27).

Such collider bias could occur through initial selection, or selective dropout, or both – for example, a study could be representative of its target population initially, but become less representative as those of poorer health drop out due to death. The main difference between these two scenarios – initial selection and selection through attrition – is in the amount of information available on the missing individuals. Where some data are available for all participants (e.g., in the case of drop-out), then multiple imputation or inverse probability weighting can be used (28), under some assumptions which are untestable given the observed data, to recover unbiased estimates of associations. However, where there is no information on missing individuals (e.g., we have no data on individuals who did not volunteer for participation into a study), then such methods cannot be used. External information (such as the expected proportion of males and females in the general population) could be used to investigate likely factors related to participation, and to derive bias-adjusted estimates.

A related issue is the use of case-control studies to examine associations with “secondary” outcomes – that is, phenotypes other than the case/control outcome (29, 30). In such studies, the association between genotype and secondary phenotype will be biased if both genotype and secondary phenotype are associated with case-control status. Case-control studies condition on case-control status, and thus again collider bias can bias the association between genotype and secondary phenotype. Various methods have been proposed to overcome this bias, including maximum likelihood and inverse probability weighting. This latter method requires some knowledge about the prevalence of case/control status in the underlying population, or the assumption that the disease is rare (29, 30).

We have discussed one important way in which selection into or out of a study can induce collider bias and spurious associations. There are other ways in which ascertainment can generate biases (31). For example, Figure 3 (panel B) shows a situation in which entry into a study is conditional upon the value of the phenotype (but not the outcome of interest) and where the phenotype does not cause the outcome, but the phenotype and outcome are correlated in unselected samples (i.e., due to genetic and/or environmental factors U). In this situation, collider bias occurs because conditioning on selection induces an association between SNPs related to the phenotype and the polygenic and/or environmental factors that influence the outcome. Therefore SNPs that cause the phenotype only (i.e. do not in truth cause the outcome), may now show spurious relationships with the outcome variable. An example of the situation in Figure 3 (panel B) is when the phenotype increases mortality (32-35) – for example, in studies of smoking as a phenotype, where smoking is associated with premature mortality. In a cohort study which examines smoking, and then follows participants up for Alzheimer’s disease, those who die early (perhaps because of smoking-related illness) will never have the chance to be diagnosed with Alzheimer’s disease, and therefore smoking will appear to be a protective factor. Figure 3 (panels C to E) also shows examples where selection will bias the estimation of the causal effects of SNPs on the outcome. In these examples, SNPs that do cause the outcome directly via the phenotype will either show increased or decreased association in the selected sample, depending on the underlying genetic and environmental aetiology of both traits. In the situations depicted in Figures 3A, 3C and 3E, the association between phenotype and outcome (e.g. in an observational study) would also be biased. In contrast, Figure 3F shows a situation where selection will bias the association of the phenotype with the outcome, but the association of the SNP with the outcome will be unbiased. Other, more complex, situations can also lead to selection bias – we have not attempted to outline every possible case here. Algorithms for deciding whether a given causal analysis is biased by selection have been described (16), and could be used to decide whether bias is likely in a given case.

**Figure 3.**
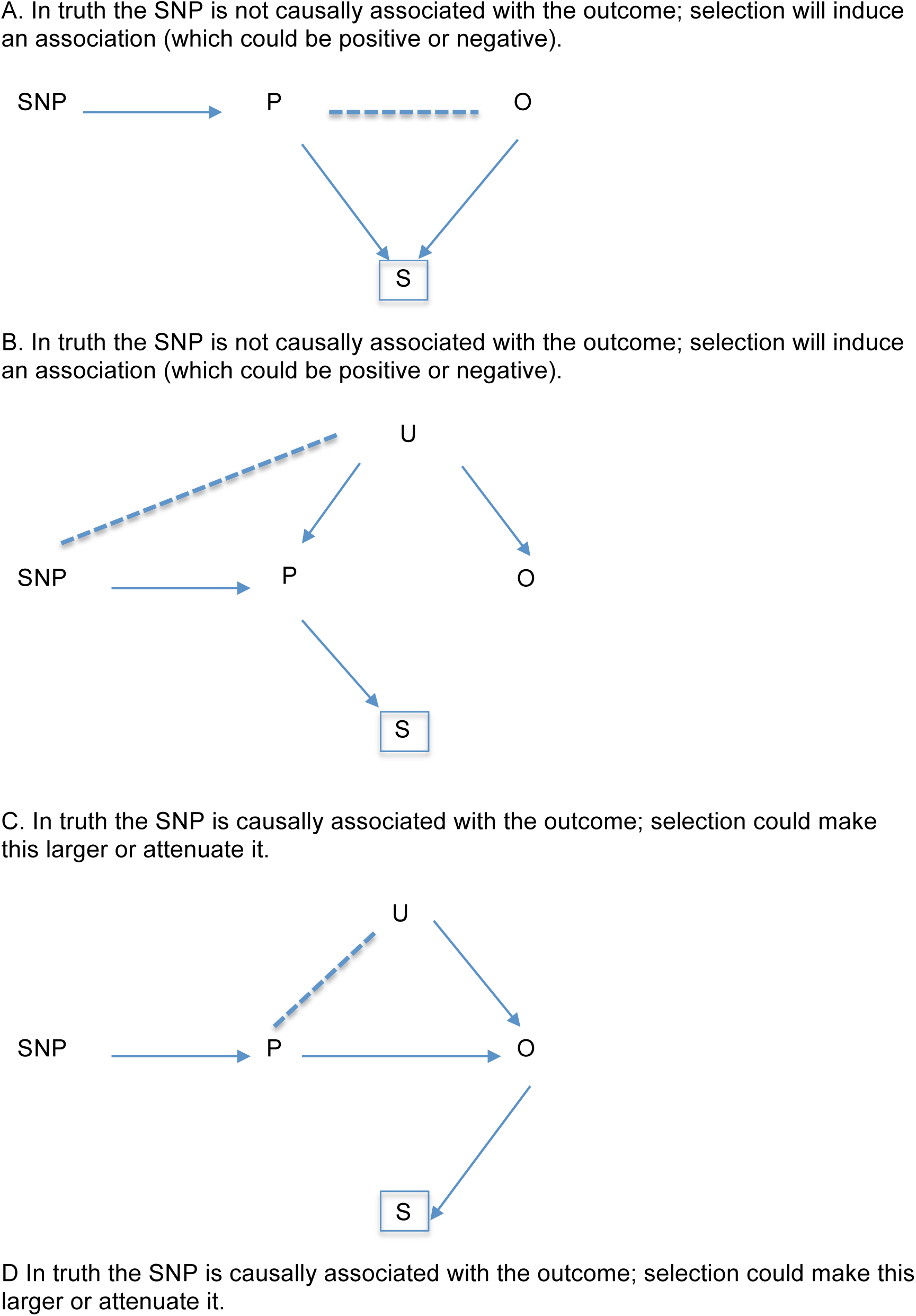
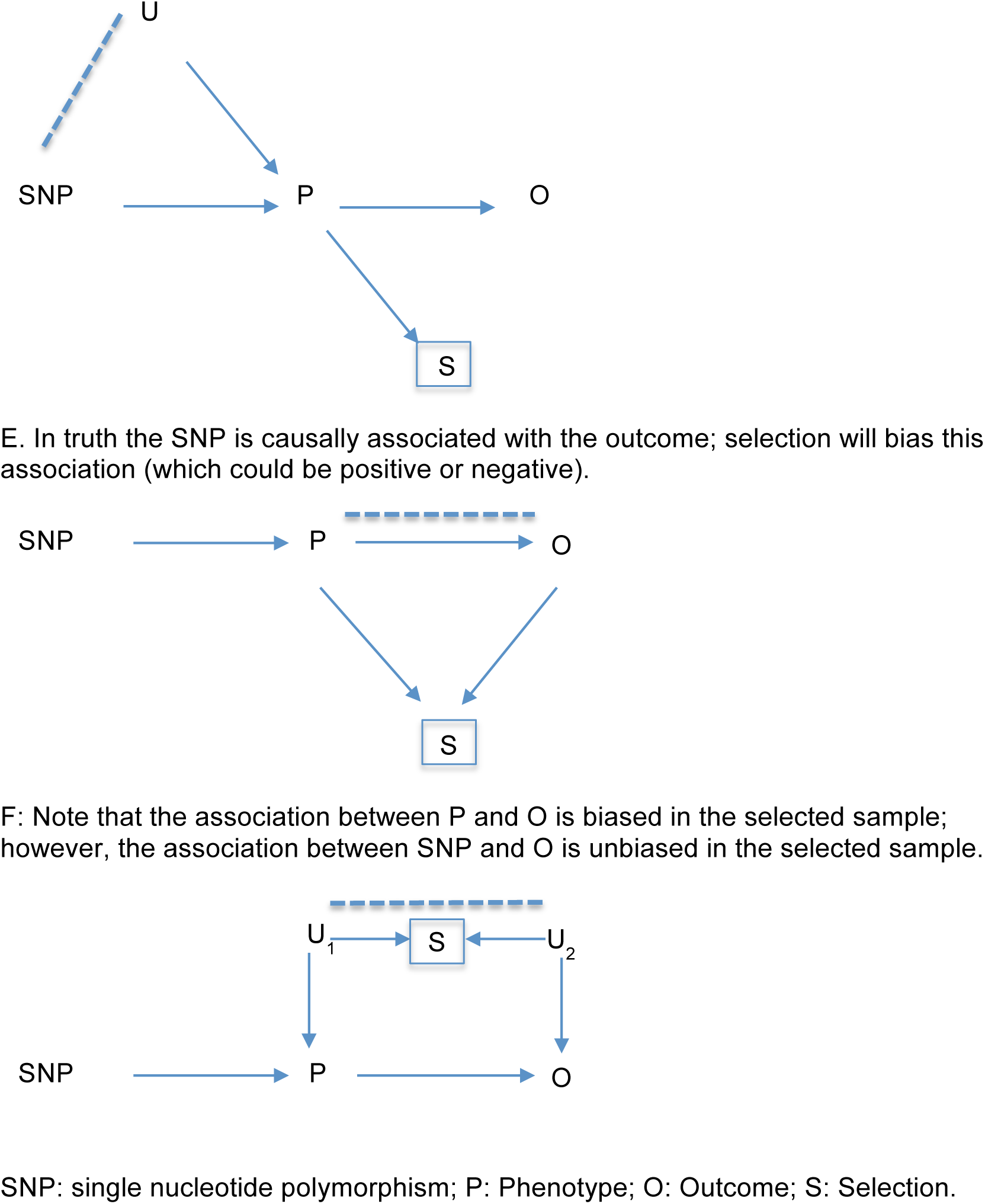
Scenarios where selection bias would occur.

Our results highlight the value of representative cohorts (including birth cohorts), where there is little or no selection into the cohort. In addition, having some baseline data and DNA available on all participants at recruitment into the study at least offers the possibility of investigating the extent to which polygenic scores (and other measured factors at baseline) predict subsequent participation. Without this knowledge, studies in samples with unknown selection/attrition mechanisms run the risk of providing biased and misleading results. In our opinion these important caveats should be borne in mind when interpreting the results of such studies.

## Acknowledgements

We are extremely grateful to all the families who took part in this study, the midwives for their help in recruiting them, and the whole ALSPAC team, which includes interviewers, computer and laboratory technicians, clerical workers, research scientists, volunteers, managers, receptionists and nurses. This publication is the work of the authors who will serve as guarantors for the contents of this paper.

